# Increased mitochondrial surface area and cristae density in the skeletal muscle of strength athletes

**DOI:** 10.1101/2023.01.15.524144

**Authors:** Javier Botella, Camilla T. Schytz, Thomas F. Pehrson, Rune Hokken, Simon Laugesen, Per Aagaard, Charlotte Suetta, Britt Christensen, Niels Ørtenblad, Joachim Nielsen

**Author notes:** **Corresponding authors:** Javier Botella and Joachim Nielsen.

## Abstract

Mitochondria are the cellular organelles responsible for resynthesising the majority of ATP. In skeletal muscle, there is an increased ATP turnover during resistance exercise to sustain the energetic demands of muscle contraction. Despite this, little is known regarding the mitochondrial characteristics of chronically strength-trained individuals and any potential pathways regulating the strength-specific mitochondrial remodelling. Here, we investigated the mitochondrial structural characteristics in skeletal muscle of strength athletes and age-matched untrained controls. The mitochondrial pool in strength athletes was characterised by increased mitochondrial cristae density, decreased mitochondrial size, and increased surface-to-volume ratio, despite similar mitochondrial volume density. We also provide a fibre-type and compartment specific assessment of mitochondria morphology in human skeletal muscle, which reveals across groups a compartment-specific influence on mitochondrial morphology that is largely independent of fibre-type. Furthermore, we show that resistance exercise leads to signs of mild mitochondrial stress, without an increase in the number of damaged mitochondria. Using publicly available transcriptomic data we show that acute resistance exercise increases the expression of markers of mitochondrial biogenesis, fission, and mitochondrial unfolded protein responses (UPR^mt^). Further, we observed an enrichment of the UPR^mt^ in the basal transcriptome of strength-trained individuals. Together, these findings show that strength athletes possess a unique mitochondrial remodelling, which minimises the space required for mitochondria. We propose that the concurrent activation of markers of mitochondrial biogenesis and mitochondrial remodelling pathways (fission and UPR^mt^) with resistance exercise may be partially responsible for the observed mitochondrial phenotype of strength athletes.

## Introduction

Mitochondria are the cellular organelles responsible for resynthesising the majority of ATP via aerobic respiration. Extensive research has shown that endurance athletes possess enhanced mitochondrial content and function (Jacobs & Lundby, 2013) and documented endurance training as a potent stimulus for enhancing mitochondrial characteristics (Holloszy, 1967; Granata *et al*., 2021) - a concept termed mitochondrial biogenesis (Botella *et al*., 2022). Conversely, strength-trained individuals have slightly increased (Staron *et al*., 1984) or similar mitochondrial content (Prince *et al*., 1981; Salvadego *et al*., 2013; Chapman *et al*., 2020) to their untrained counterparts despite increased function (Salvadego *et al*., 2013). However, the effect of resistance training on mitochondrial adaptations remains equivocal (Parry *et al*., 2020) and only sparsely examined. Previous research has shown that 6 months of resistance training led to decreased citrate synthase (CS) activity (Tesch *et al*., 1987) (a marker of mitochondrial content (Larsen *et al*., 2012)). Subsequent studies have shown decreased (MacDougall *et al*., 1979; Roberts *et al*., 2018), or unchanged (Tesch *et al*., 1990; Salvadego *et al*., 2013; Porter *et al*., 2015; Flack *et al*., 2016; Groennebaek *et al*., 2018) markers of mitochondrial content following resistance training; with only a single study showing an increase (Tang *et al*., 2006). In contrast, mitochondrial respiratory function has been shown to either increase (Salvadego *et al*., 2013; Porter *et al*., 2015; Groennebaek *et al*., 2018; Holloway *et al*., 2018) or remain unchanged (Flack *et al*., 2016; Robinson *et al*., 2017), but never to decrease in response to resistance training. Collectively, these findings would seem to suggest that non-volumetric mitochondrial remodelling could underlie the specific mitochondrial plasticity observed with resistance training. It has been suggested that the increases in fibre cross-sectional area (fCSA) observed following resistance training may dilute, and therefore mask, the magnitude of mitochondrial volumetric adaptations (MacDougall *et al*., 1979; Lüthi *et al*., 1986; Parry *et al*., 2020). Therefore, given that the use of biochemical assays to interrogate mitochondrial adaptations (Larsen *et al*., 2012; Kuang *et al*., 2022) are unable to account for difference in fCSA, the use of transmission electron microscopy (TEM) may better reveal the distinct structural mitochondrial characteristics of resistance-trained individuals, such as mitochondrial morphology (Castro-Sepulveda *et al*., 2020) and cristae density (Nielsen *et al*., 2017b), that are independent of fCSA.

Phenotypical adaptations following months or years of training stem from the accumulative effect of repeated exercise over time (Perry *et al*., 2010; Viggars *et al*., 2023). The acute molecular cascade initiated following an acute resistance exercise bout involves the upregulation of a large number of genes including ribosomal and myogenic pathways that will support the increased synthesis of myofibrillar proteins (Bamman *et al*., 2018). In addition, resistance exercise upregulates the expression of PPARG coactivator 1 alpha (PGC1α) (Ydfors *et al*., 2013) - the so called master regulator of mitochondrial biogenesis. However, an alternative splicing pattern of PGC1α appears to be elicited by resistance exercise (Ydfors *et al*., 2013; Silvennoinen *et al*., 2015) and to divergently contribute to the phenotypical adaptation to resistance exercise (Ruas *et al*., 2012). However, little is known regarding the effects of resistance exercise *per se* on mitochondrial structural disturbance, and other signalling, that could subsequently affect any mitochondrial-related remodelling.

The aim of the present study, therefore, was to elucidate the mitochondrial morphological and structural characteristics of chronically trained strength athletes compared to untrained age-matched individuals. To achieve this, we extensively characterised the compartment- and fibre-type-dependent mitochondrial morphology of the human *vastus lateralis* muscle. Furthermore, we aimed to decipher the mitochondrial morphological changes following acute resistance exercise and, using publicly available datasets, the mitochondrial-related transcriptional signalling in response to resistance exercise and at rest in strength-trained individuals.

## Material & Methods

### Ethical approval

The human skeletal muscle biopsy material included in the present analysis originates from two previous published reports (Nielsen *et al*., 2017a; Hokken *et al*., 2021). The participants were fully informed of potential risks associated with the experimental conditions before obtaining their verbal and written consent. The project with strength athletes was approved by the local Ethics Committee in the Region of Southern Denmark (project ID S-20160116) and the study with untrained males was approved by the Local Human Ethical Committee of the Central Denmark Region (M-20110035). The experiments conformed to the standards set by the Declaration of Helsinki.

### Participants

Participants’ biological and anthropometric characteristics are summarised in Table 1. Biopsy samples were obtained from the *vastus lateralis* muscle in ten male elite power lifters before and after (< 5 min) an acute resistance exercise session, and in a resting condition of twelve age-matched untrained males (Table 1; Figure 1.A and Figure 5.A).

**Table 1.** Biological and anthropometric participant characteristics. NA, not available. RM, one-repetition maximum. The 1 RM was self-reported. * = significantly different from untrained (*P* < 0.05). Values are mean ± SD.

| | n | Age<br>(y) | Weight<br>(kg) | $\dot{V}\text{O}_{2\text{max}}$<br>(ml min<sup>-1</sup> kg<sup>-1</sup>) | 1RM back squat<br>(kg) | Reference |
| --- | --- | --- | --- | --- | --- | --- |
| Untrained | 12 | 23 ± 5 | 76 ± 8 | 41 ± 7 | NA | (Nielsen *et al.*, 2017a) |
| Strength athletes | 10 | 24 ± 4 | 95 ± 8 * | NA | 210 ± 25 | (Hokken *et al.*, 2021) |

**Figure 1.**
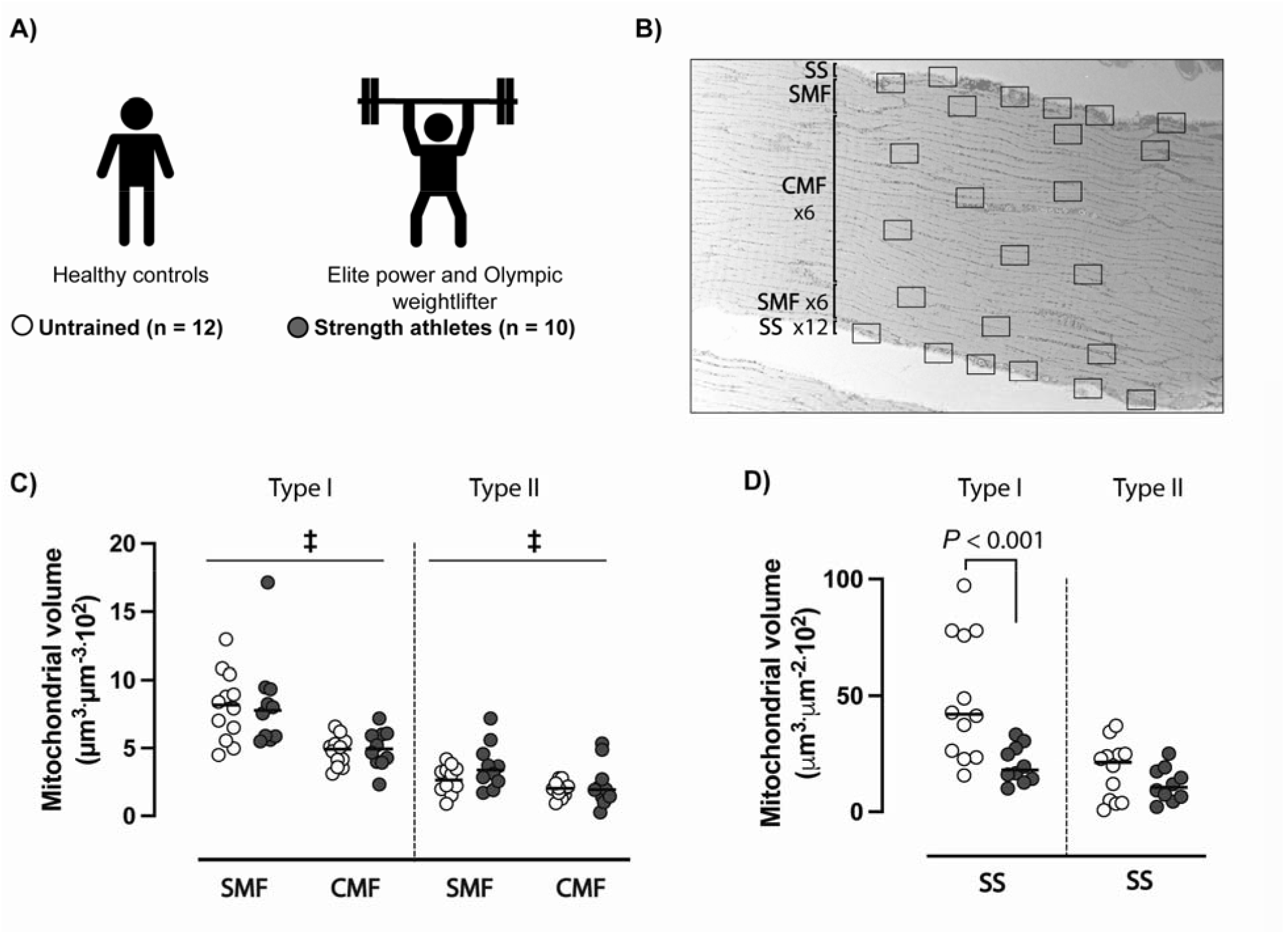
Skeletal muscle mitochondrial content among untrained individuals and strength athletes. A) Description and sample size of the groups studied in the study. B) Transmission electron microscopy (TEM) micrograph defining the distinct compartments within skeletal muscle. C) Mitochondrial volumetric density (V_V_) of superficial and central intermyofibrillar; and D) mitochondrial volumetric density (V_A_) of subsarcolemmal mitochondria in type I and II skeletal muscle of untrained individuals (n = 12) and strength athletes (n = 10). ‡ = compartment effect (*P* < 0.05).

### Maximal oxygen uptake (V□O_2_max)

Maximal oxygen uptake of the untrained individuals was measured on an ergometer bicycle (Monark Ergomedic 828E; Monark, Varberg, Sweden). The test was performed at a constant pedaling rate of 70 rpm, and the first 5 min at 140 Watt served as a warm-up to reach a steady state of submaximal work. Hereafter, the workload was increased with 35 Watt every minute until exhaustion. The oxygen uptake was measured every 10 s (AMIS 2001; Innovision, Odense, Denmark), and V□O_2_max was calculated as the highest mean of three consecutive measurements.

### Resistance exercise protocol

The exercise protocol is described in detail elsewhere (Hokken *et al*., 2021). In brief, the session consisted of three lower body exercises (barbell back squats, barbell deadlifts from a deficit and dumbbell rear foot elevated split squats) and performed with training loads of 40%-75% of subject’s self-reported 1 repetition maximum (1RM). Subjects performed three warm-up sets in the barbell back squat consisting of 10 repetitions at 40%, 8 repetitions at 50% and 6 repetitions at 60% of 1RM. Subsequently, four sets were performed consisting of 5 repetitions at 70%-75% 1RM. One warm-up set of 5 repetitions at 60% 1RM was then completed in the barbell deadlift exercise, followed by 4 sets of 5 repetitions at 70-75% 1RM. Lastly, 4 sets of 10-12 repetitions in the split squat exercise were performed on each leg, alternating the working leg between successive sets. Rest intervals between sets were maintained at 3-6 minutes in the barbell back squat and the barbell deadlift, while using 1-2 minute rest periods in the split squats.

### Transmission electron microscopy

Muscle specimens were prepared for TEM as described in detail elsewhere (Jensen *et al*., 2022). Upon collection, specimens were immediately fixed with a 2.5% glutaraldehyde in 0.1 mol/L sodium cacodylate buffer (pH 7.3) for 24 hours at 4°C and subsequently rinsed four times in 0.1 mol/L sodium cacodylate buffer. Following rinsing, fibres were post-fixed with 1% osmium tetroxide (OsO4) and 1.5% potassium ferrocyanide (K4Fe(CN)6) in 0.1 mol/L sodium cacodylate buffer for 90 minutes at 4°C. After post-fixation, the fibres were rinsed twice in 0.1 mol/L sodium cacodylate buffer at 4°C, dehydrated through a graded series of alcohol at 4-20°C, infiltrated with graded mixtures of propylene oxide and Epon at 20°C, and embedded in 100% Epon at 30°C. Ultrathin (60 nm) sections were cut (using a Leica Ultracut UCT ultramicrotome) in three depths (separated by 150 μm) and contrasted with uranyl acetate and lead citrate. Sections were examined and photographed in a pre-calibrated transmission electron microscope (JEM-1400Plus, JEOL Ltd, Tokyo, Japan and a Quemesa camera, or Philips EM 208; Philips, Amsterdam, The Netherlands, and a Megaview III FW camera, Olympus; Soft Imaging Solutions, Münster, Germany). All longitudinally oriented fibres (n = 6-12) were photographed at ×10-13,000 magnification in a randomized systematic order, including 12 images from the subsarcolemmal region and 12 images from the myofibrillar region (Figure 1.B). The individual fibres were classified as either type 1 or 2 based on their intermyofibrillar mitochondrial volume fraction and Z-line width as previously described (Sjöström *et al*., 1982) and modified using the present TEM protocol (Jensen *et al*., 2022).

### Mitochondrial volume and cristae density

The mitochondrial content of the myofibres was assessed by point counting (Weibel, 1980) using a grid size of 135 nm. Central and superficial intermyofibrillar mitochondria (CMF and SMF, respectively) were expressed per myofibrillar space (µm^3^ µm^-3^) and subsarcolemmal (SS) mitochondrial content was expressed relative to the surface area of the outermost myofibril (µm^3^ µm^-2^). Mitochondrial cristae density was assessed by counting intersections of cristae on test lines (*I*_L_) multiplied by 2, where *I* is the number of intersections and L is the total length of test lines using a grid size of 270 nm (Nielsen *et al*., 2017b).

### Mitochondrial morphology

Mitochondria were manually traced, and predefined shape descriptors were obtained using ImageJ (v.1.53a, NIH, http://rsb.info.nih.gov/ij) as previously reported (Picard *et al*., 2013). Mitochondrial morphological variables were studied from the three different mitochondrial compartments (SS, SMF, CMF) according to previous classifications (Hokken *et al*., 2021). All mitochondrial analyses were performed blinded to participant id and timepoint. In total, 31,454 individual mitochondria were manually traced including 14,138 mitochondria from the untrained individuals (SS = 3,608; SMF = 5,534; CMF = 4,996), while 17,316 mitochondria were identified from the strength athletes at rest (SS = 2,182; SMF = 4,096; CMF = 1,960) and following exercise (SS = 2,785; SMF = 4,390; CMF = 1,903). Illustration of mitochondrial morphology distribution was performed using R studio with the ‘geom_density’ function of *ggplot2*.

### Mitochondrial integrity

Mitochondrial integrity was evaluated as ‘healthy’ or ‘damaged’ in the resting and post-resistance exercise micrographs obtained from the strength athletes. Mitochondria were defined as ‘healthy’ if the cristae were able to span across opposite sides of the outer membrane and preserved a high electron density (e.g., dark colour; Figure 5.E). Mitochondria were labelled as ‘damaged’ when the following indicators were observed: 1) reduced electron density within the mitochondrion (e.g., reduced darkness of inner mitochondrial sections) suggestive of permeabilised outer membrane, and 2) decreased cristae numbers with some cristae unable to span across opposite sides of the outer membrane (see red arrows in Figure 5.E). These different categories were established based on the mitochondrial cristae dynamics observed following induction of *in vitro* mitochondrial stress (Wang *et al*., 2019). All analyses were performed in a randomised order blinded from participant id and timepoint information.

### Acute regulation of gene expression following resistance exercise

To obtain gene expression data from human skeletal muscle in response to resistance exercise, we used the publicly available database MetaMEx (Pillon *et al*., 2020) that combines all published and available transcriptomic datasets in human studies. We selected ‘resistance exercise studies’ and restricted the analyses to 0 to 6 hours post-exercise, without any other participant criteria, to obtain a high statistical power and a large sample size (n = 148). Genes from each of the pathways of interest (Figure 6.A) were manually curated based on previous literature (Pakos-Zebrucka *et al*., 2016; Trewin *et al*., 2018; Bishop *et al*., 2019; Huertas *et al*., 2019; Guo *et al*., 2022).

### Gene expression differences between strength-trained and untrained individuals

Skeletal muscle RNA-sequencing data from a published study (Chapman *et al*., 2020) was reanalysed to establish the gene expression profile of strength-trained (n = 7 males) and untrained (n = 7 males) individuals. In brief, following the download of the raw fastq files, gene read counts were obtained using Kallisto (Bray *et al*., 2016), count data was normalised in R using *edgeR*, and differential expression analysis was performed in R using *limma* (Law et al., 2018). Gene-set enrichment analyses (GSEA) was performed using the R package fast geneset enrichment analysis (*FGSEA*) (Korotkevich *et al*., 2021). Mitochondrial-specific genesets were utilised (obtained from the MitoCarta 3.0 (Rath *et al*., 2021)) and the mitochondrial unfolded protein response (UPR^mt^) geneset was obtained from a mouse study (Guo *et al*., 2022). An adjusted p-value lower than 0.05 was deemed significant and enrichment values are reported as normalised enrichment scores (NES).

### Statistical analysis

Statistical analysis was performed using Stata, version 17 (StataCorp LP). All interactions or main effects were tested using a linear mixed-effects model, with group, type, compartment, and time as fixed effects, and subject as random effect. Data assumptions on heteroscedasticity and normal distribution were evaluated by inspecting the distribution of residuals and a standardized normal probability plot, respectively. If necessary, data were transformed prior to analysis. The difference between fiber types and compartments, and the effects of acute exercise on mitochondrial shape descriptors were conducted based on the participants median values. Unless otherwise stated, values are presented as median ± standard deviation. Significance level was set at *p* < 0.05.

## Results

### Mitochondrial volume density

Mitochondrial volumetric density was not different across untrained individuals and strength athletes across fibre types and in both CMF and SMF, averaging in CMF (4.9 ± 1.1 vs 4.9 ± 1.4 µm^3^ µm^-3^ 10^2^ for type I fibres of untrained individuals and strength athletes, respectively; 2.0 ± 0.6 vs 1.9 ± 1.6 µm^3^ µm^-3^ 10^2^, respectively, for type II fibres; Figure 1.C) and SMF (8.2 ± 2.5 vs 7.8 ± 3.5 µm^3^ µm^-3^ 10^2^, respectively, for type I fibres; 2.7 ± 1.0 vs 3.4 ± 1.7 µm^3^ µm^-3^ 10^2^, respectively, for type II fibres; Figure 1.C). There was a significant difference in the abundance of SS mitochondria of strength athletes, compared to controls, in type I fibres (42.2 ± 26.8 vs 18.1 ± 8.0 µm^3^ µm^-2^ 10^2^, respectively; Figure 1.D), but not in type II fibres (21.4 ± 12.5 vs 10.6 ± 7.3 µm^3^ µm^-2^ 10^2^, respectively; Figure 1.D).

### Skeletal muscle mitochondrial morphology

Despite similar mitochondrial volumetric density, strength athletes possessed a larger number of mitochondrial profiles (numerical density of mitochondria; # µm^-3^ or # µm^-2^) in the myofibrillar compartments of both type I and II fibres compared to controls (Figure 2.A). This disparity was not observed for SS mitochondria (Figure 2.B). Measures of mitochondrial morphology showed that strength athletes had a smaller mitochondrial size (area) across all intracellular compartments and in both type I and II fibres (Figure 2.C), along with a reduced circularity (Figure 2.D), but comparable aspect ratio (Figure 2.E). Together, the observed group differences led to a larger surface-to-volume ratio per mitochondrion in the strength athletes vs controls (Figure 2.F). Of interest, across groups and compartments, type I fibres demonstrated largely similar mitochondrial morphology as in type II fibres (Figure 4.A), indicating that the mitochondrial volumetric difference observed between type I and II fibres was mainly due to an increased number of mitochondrial profiles (Figure 4.C and 4.D). Lastly, across groups and fibre types, SS and SMF mitochondria were characterised by large area, lower circularity, and increased aspect ratios when compared to CMF mitochondria (Figure 4.B).

**Figure 2.**
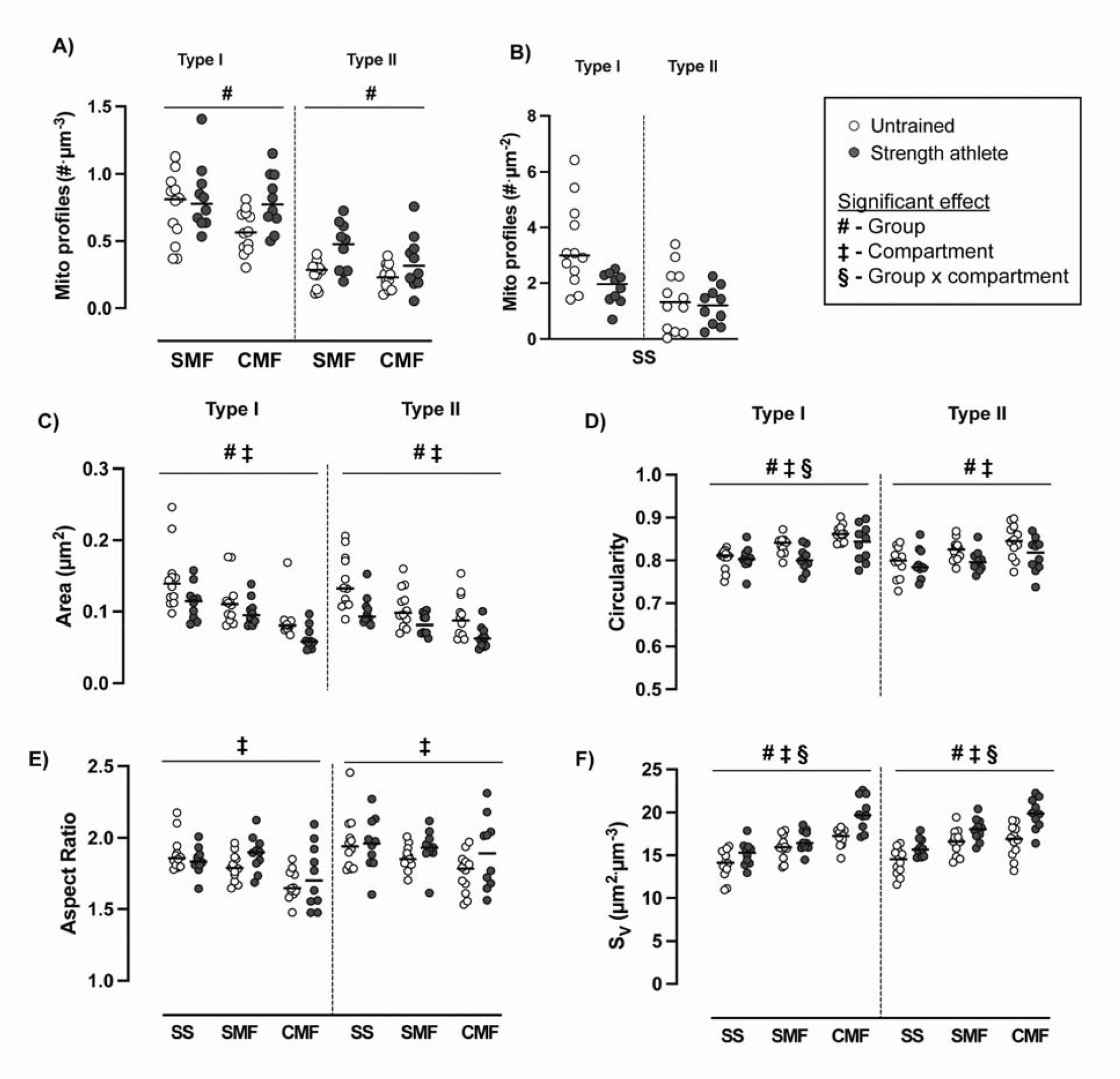
Morphological characteristics of skeletal muscle mitochondria in untrained individuals and strength athletes. A) Mitochondrial profiles in intermyofibrillar; and B) subsarcolemmal compartments across fibre types. C) Mitochondrial area; D) mitochondrial circularity; E) mitochondrial aspect ratio; and F) mitochondrial surface to volume ratio (S_V_) across fibre types and compartments in untrained individuals (n = 12) and strength athletes (n = 10). # = group effect (*P* < 0.05); ‡ = compartment effect (*P* < 0.05); § = group and compartment effect (*P* < 0.05). SS = subsarcolemmal; SMF = superficial intermyofibrillar; CMF = central intermyofibrillar.

### Mitochondrial cristae density

Given the distinct mitochondrial morphology of strength athletes, the density of mitochondrial cristae was evaluated in a subset of participants that had sufficient mitochondria with clearly defined cristae structures (≥8 mitochondria to obtain a reproducible value) in untrained individuals (n = 9) and strength athletes (n = 5). We observed a 16% increase in the mitochondrial cristae density of strength athletes when compared to untrained (30.9 ± 1.8 vs 26.6 ± 2.4 µm^2^ µm^-3^; Figure 3.A), with strength athletes demonstrating values comparable to those previously observed in highly trained endurance athletes (Nielsen *et al*., 2017b).

**Figure 3.**
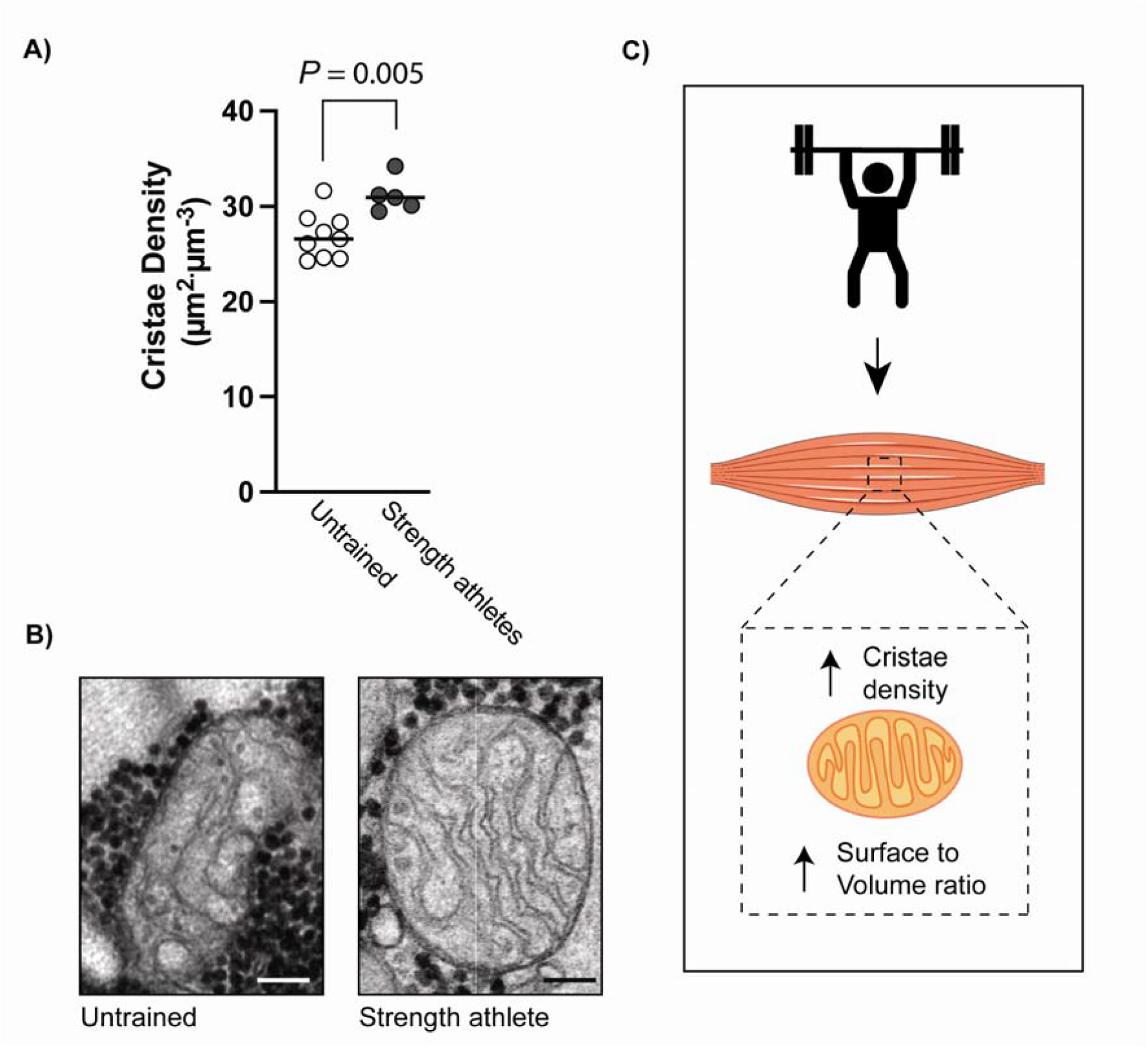
Mitochondrial cristae density in untrained individuals and strength athletes. Mitochondrial cristae density in untrained individuals (n = 9) and strength athletes (n = 5). Representative micrograph showing a mitochondrion from an untrained individual and a strength athlete. C) Schematic illustration of the mitochondrial structural differences between groups. Scale bar = 0.1 µm.

**Figure 4.**
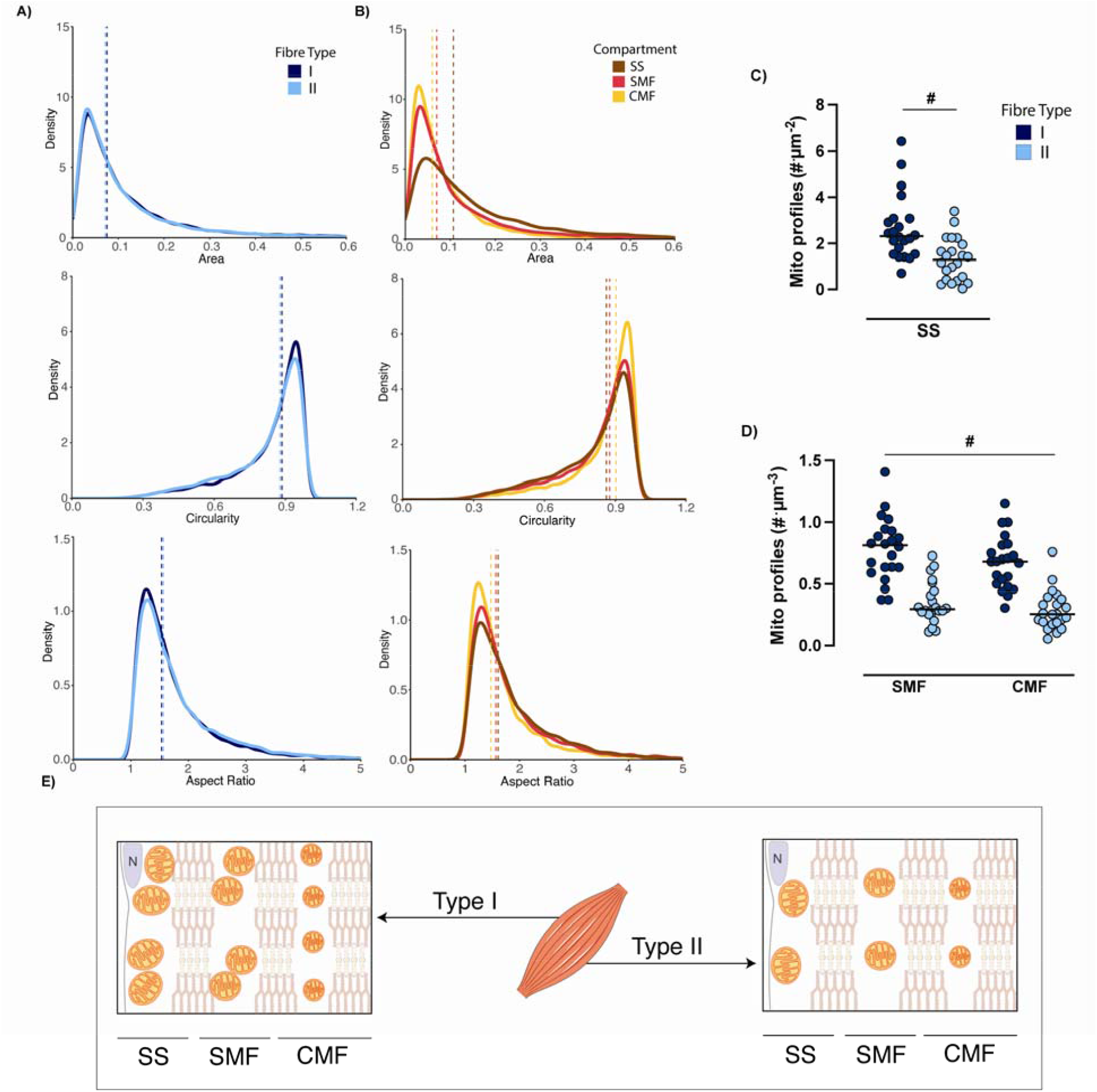
Mitochondrial morphological characteristics of human skeletal muscle. A) Density distribution of mitochondrial area, circularity, and aspect ratio between type I (n = 13,154 mitochondria) and type II (n = 9,220 mitochondria) fibre types. B) Density distribution of mitochondrial area, circularity, and aspect ratio between subsarcolemmal (SS; n = 5,769 mitochondria), superficial intermyofibrillar (SMF; n = 9,649 mitochondria), and central intermyofibrillar (CMF; n = 6,956 mitochondria) skeletal muscle fibres. C) Mitochondrial profile number in the subsarcolemmal; and D) in the intermyofibrillar regions across fibre types. E) Schematic model of the observed morphological and numerical density differences observed between type I and II fibres, as well as across fibre compartments. Type I fibre = dark blue; type II fibre = light blue; SS mitochondria = brown; SMF mitochondria = red; CMF mitochondria = yellow. # = fibre type effect (*P* < 0.05).

### Effects of an acute resistance exercise session in strength athletes

The mitochondrial structural and network differences presently observed between strength athletes and untrained individuals may suggest that acute resistance exercise could activate mitochondrial-related pathways, in turn leading to the observed mitochondrial phenotypical remodelling in strength athletes. Exercise-induced mitochondrial morphological stress has been shown to be influenced by the specific exercise protocol (Picard *et al*., 2013; Huertas *et al*., 2019), but little is known about the induction of mitochondria with acute resistance exercise. Using TEM micrographs obtained before and immediately after a resistance exercise session (Figure 5.A), marked changes in the SS mitochondrial network were observed as manifested by larger profile areas, lower aspect ratios, and greater circularity, which together resulted in a reduced surface-to-volume ratio (Figure 5.B, 5.C and 5.D) These effects were smaller or absent in SMF and CMF mitochondria (data not shown) and did not differ between fibre types (*P* = 0.20 – 0.94). Next, mitochondrial structural changes, such as electron density or cristae integrity, were used as a proxy of mitochondrial integrity (Figure 5.E). Despite increased signs of mitochondrial damage following exercise in some individuals, there was no systematic exercise effect in either type I (SS: 0 ± 1 to 0 ± 5%; SMF: 0 ± 1 to 0 ± 6%; CMF: 0 ± 0 to 0 ± 5%; *P* = 0.77) or type II fibres (SS: 0 ± 2 to 2 ± 24%; SMF: 0 ± 1 to 1 ± 23%; CMF: 0 ± 1 to 0 ± 24%; *P* = 0.17). However, those mitochondria identified as damaged (Figure 5.E) were significantly larger in area when compared to non-damaged mitochondria, indicative of mitochondrial swelling (*P* < 0.05; Figure 5.F).

**Figure 5.**
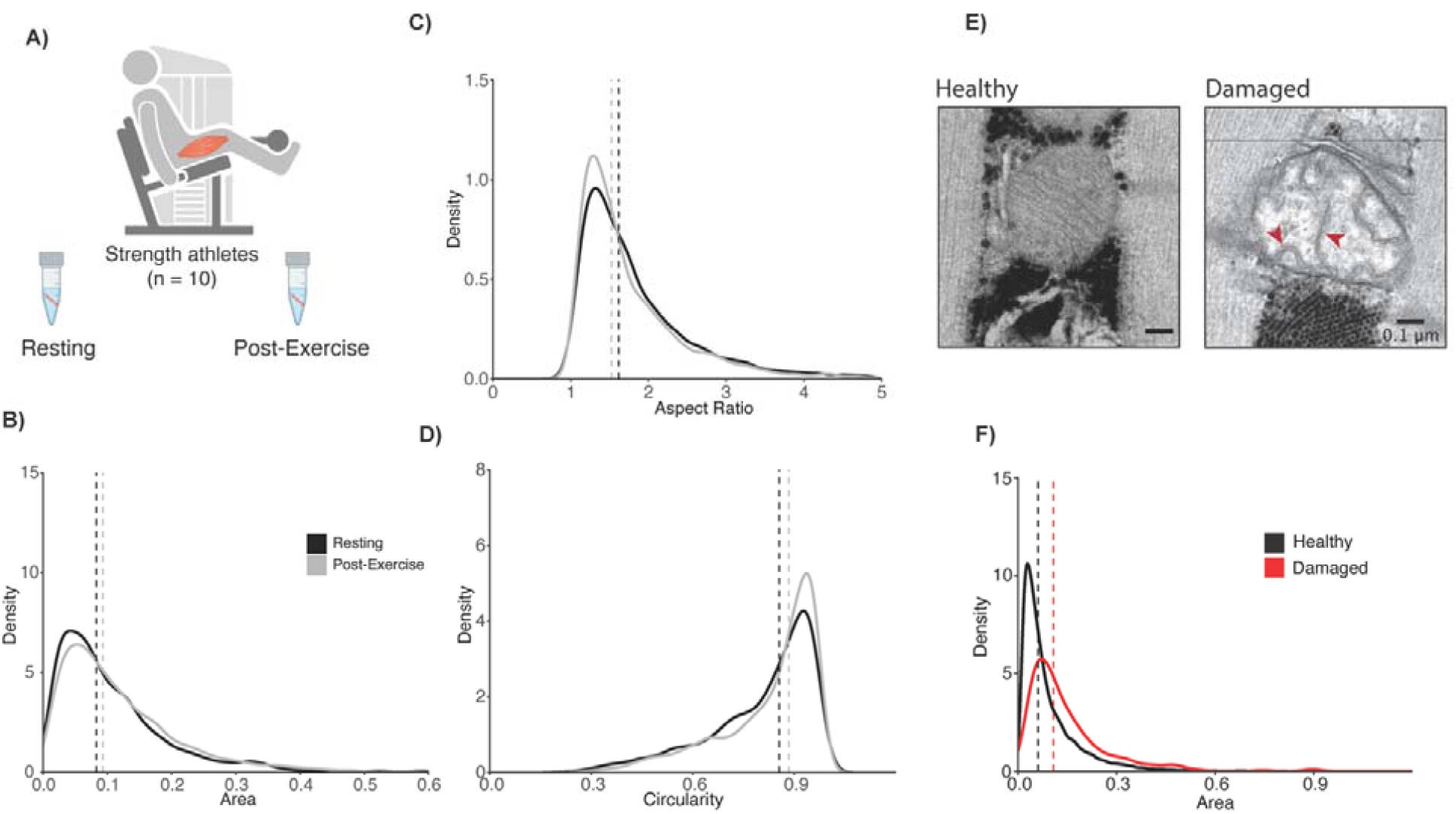
Mitochondrial morphological changes in skeletal muscle following a resistance exercise session in strength athletes. A) Schematic representation of the muscle biopsies obtained before and after resistance exercise in strength athletes (n = 10). B) Changes in the mitochondrial area; C) circularity; and D) aspect ratio in subsarcolemmal mitochondria (SS; resting = 2,182 mitochondria, post-exercise = 2,785 mitochondria). E) Representative image of a healthy and damaged mitochondria, with red arrows pointing at cristae abnormalities; and F) density distribution of mitochondrial area between healthy and damaged mitochondria. Resting sample = black; post-exercise sample = grey.

### Gene expression following acute resistance exercise and in strength-trained individuals

We next sought to interrogate mitochondrial-related transcriptional pathways, such as mitochondrial biogenesis and dynamics, following acute resistance exercise. We observed that genes involved in the regulation of mitochondrial biogenesis were upregulated acutely following resistance exercise (Figure 6.A). Conversely, most of the genes involved in the different branches of mitochondrial dynamics were not upregulated. One exception was *MIEF2*, a known mitochondrial receptor that recruits mitochondrial fission factor (*MFF*) to the fission site (Osellame *et al*., 2016). Another pathway that is involved in mitochondrial remodelling is the mitochondrial unfolded protein response (UPR^mt^), which is involved in mitochondrial protein homeostasis (Mottis *et al*., 2019). Two of the main mitochondrial proteases, *HSPD1* and *HSPE1*, were upregulated following resistance exercise. In mammals, concurrently to UPR^mt^ is the activation of the integrated stress response (ISR) (Pakos-Zebrucka *et al*., 2016), with two of its major players, *ATF4* and *DDIT3* also found to be upregulated following acute resistance exercise (Figure 6.A). Similarly, the negative regulators of eukaryotic initiation factor (eIF2α) phosphorylation (primary sensors of ISR activation) (Pakos-Zebrucka *et al*., 2016), *PPP1R15A* and *PPP1R15B*, also seemed to be upregulated suggesting an activation of the ISR response (Figure 6.A). Last, we aimed to establish the gene expression signatures of strength-trained individuals when compared to untrained counterparts using a previously published dataset (Chapman *et al*., 2020). We observed an enrichment of the UPR^mt^ geneset (adjusted *p* = 0.02; NES = 1.35; Figure 6.B), but not of other mitochondrial genesets such as mitochondrial autophagy (adjusted *p* = 1; NES = 0.44) or mitochondrial dynamics (adjusted *p* = 1; NES = 0.86). These results suggest that UPR^mt^ is a unique signature of chronically strength-trained individuals and may play a role in the mitochondrial phenotypical remodelling observed in these athletes (Figure 6.C).

**Figure 6.**
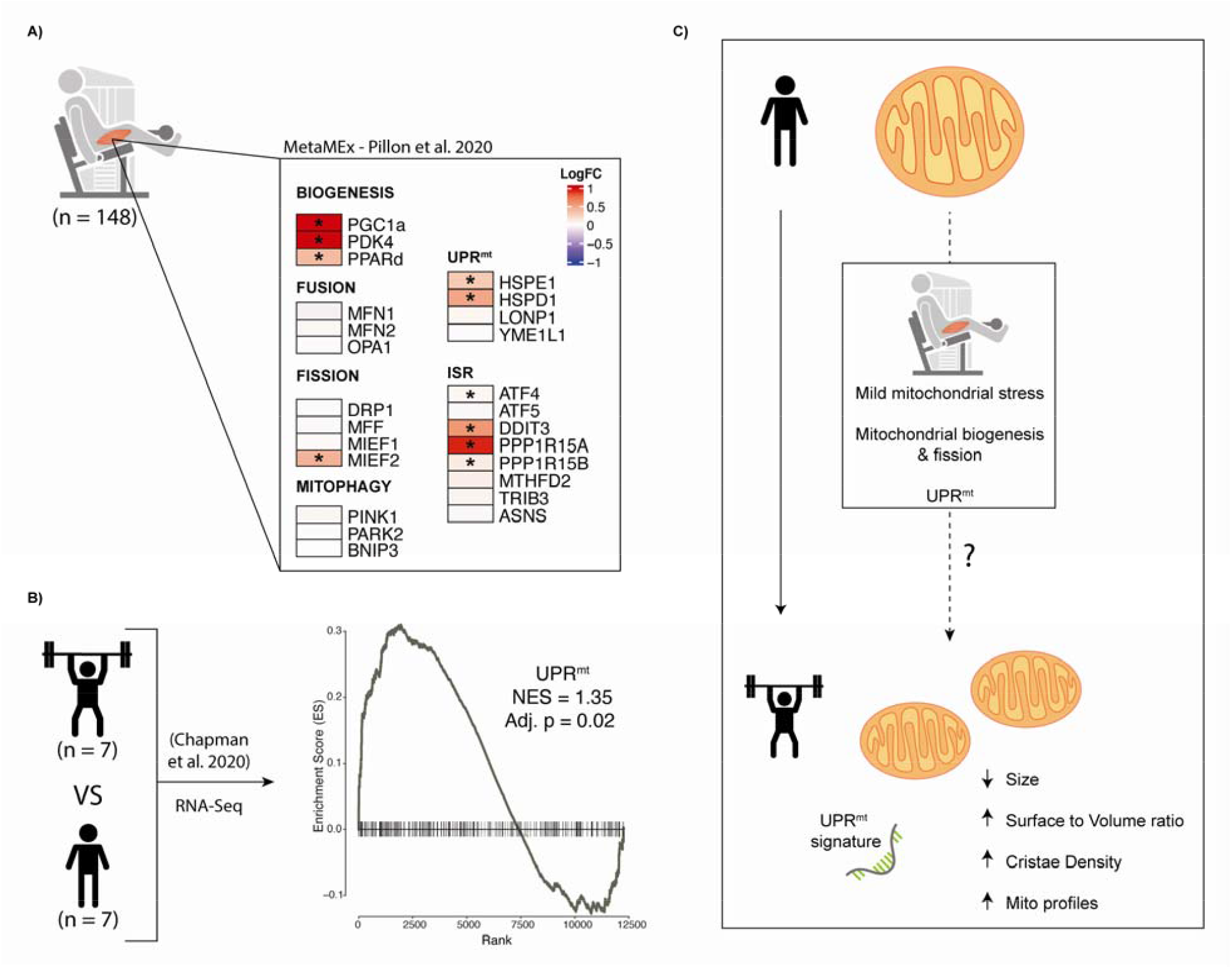
Transcriptional response to resistance exercise and hypothesised model of mitochondrial remodelling. A) Transcriptional changes, in genes involved in mitochondrial-related pathways (mitochondrial biogenesis, fusion, fission, mitophagy and mitochondrial unfolded protein response [UPR^mt^]) and the integrated stress response (ISR), following resistance exercise in human skeletal muscle (0 to 6 hours; n = 148) (Pillon *et al*., 2020); B) Gene-set enrichment analysis (GSEA) of the UPR^mt^ geneset between untrained and strength-trained individuals (Chapman et al., 2020); C) Proposed model by which chronic resistance exercise may lead to phenotypical changes in the mitochondrial remodelling of human skeletal muscle.

## Discussion

The present data reveal that substantial mitochondrial structural remodelling seems to occur in skeletal muscle of chronically strength-trained athletes (elite power and Olympic weightlifters). Despite similar mitochondrial volumetric density, the mitochondria of strength athletes were characterised by smaller size, increased number of mitochondrial profiles, increased cristae density, and increased surface-to-volume ratio - which may represent an important adaptive response to the high metabolic demands of resistance exercise. We also report that human skeletal muscle fibre types I and II differ in the numerical density of mitochondria profiles (i.e., showing differently expanded networks), which leads to volumetric differences, rather than disparities in morphological profiles. Our results also show distinct mitochondrial morphology across the myofiber space, with bigger and more complex mitochondria situated in the subsarcolemmal compartments, which decreases with myofiber depth. Furthermore, acute resistance exercise in trained individuals seems to result in compartment-specific increases in morphological indicators of mitochondrial stress, such as increased area and circularity in SS mitochondria, however, without resulting in more damaged mitochondria. This observation of mild resistance-exercise induced mitochondrial stress is further supported by the increased transcription of genes involved in the regulation of mitochondrial biogenesis, fission, and the UPR^mt^ - the latter representing a unique signature of strength-trained individuals.

### Mitochondrial characteristics in strength athletes

Our findings suggest that chronically resistance-trained individuals possess a different skeletal muscle mitochondrial structure and network compared to untrained individuals. However, we observed no differences in mitochondrial volume density between untrained individuals and strength athletes, except for the subsarcolemmal (SS) mitochondria of type I fibres, which were more abundant in untrained individuals. These results are in line with previous reports showing no differences in markers of mitochondrial content between untrained and resistance trained individuals (Prince *et al*., 1981; Salvadego *et al*., 2013; Chapman *et al*., 2020) when evaluated at the whole muscle level. The present observation of decreased mitochondrial volume in type I SS mitochondria of chronically strength trained athletes could explain the reported reduced oxygen extraction capacity of these individuals (Salvadego *et al*., 2013), which would be especially important in mitochondria-abundant type I fibres. To the best of our knowledge, this is the first time that mitochondrial characteristics, other than volume density (Prince *et al*., 1981; MacDougall *et al*., 1982; Staron *et al*., 1984), have been extensively studied using TEM micrograph analysis in strength athletes. Our analysis revealed that the mitochondrial network of strength athletes, when compared to controls, can be characterised by smaller mitochondria, despite similar mitochondrial volumetric density, which translates into increased numerical density of mitochondrial profiles and a larger surface-to-volume ratio. Previous research have shown greater mitochondrial surface-to-volume ratio in mouse glycolytic fibres, when compared to oxidative fibres, allowing mitochondria to better interact with their surrounding environment (Bleck *et al*., 2018). This may indicate that the mitochondria of strength athletes may be better fitted to handle conditions of fluctuating and high energy turnover - as required during heavy-load and/or explosive-type resistance exercise. We hypothesise that the present signs of mitochondrial remodelling is the result of the accumulated years of resistance training, as a previous study showed no surface-to-volume ratio change following 6 weeks of heavy resistance training (Lüthi *et al*., 1986). The observed mitochondrial adaptation may also be specific to resistance training as 6 weeks of endurance training did not result in an increased number of mitochondrial profiles (Meinild Lundby *et al*., 2018).

Our analyses showed that strength athletes possess an increased mitochondrial cristae density. Increased cristae density in human skeletal muscle was first described in highly trained endurance athletes and professional soccer players (Nielsen *et al*., 2017b), and was suggested to occur in muscles of highly trained individuals, where there is a large competition for intramuscular space, and therefore the need to increase mitochondrial-specific respiratory function without increasing volume. Given the importance of the muscle contractile machinery in muscle strength and power, it is not surprising that cristae density adaptations have occurred in this population to minimise the space required for mitochondria. Combined, our results may provide an explanation for the increased mitochondrial-specific respiratory function previously reported in trained individuals (Salvadego *et al*., 2013) and observed following resistance training (Porter *et al*., 2015; Groennebaek *et al*., 2018). Whether these adaptations may influence strength performance metrics (i.e., power), or are associated with the protective role of resistance training in metabolic health and lifespan, requires further research. Similarly, future research should investigate whether the benefits of adding resistance training to an endurance training program (Aagaard & Andersen, 2010; Aagaard *et al*., 2011) may partially be due to optimised mitochondrial structure through morphological remodelling and increased cristae density.

### Fibre-type and compartment specific mitochondrial morphology

Increased mitochondrial volumetric density is a characteristic of human type I fibres (Sjöström *et al*., 1982; Reisman *et al*., 2022) and of mouse oxidative fibres (Glancy *et al*., 2017; Bleck *et al*., 2018). Our results indicate that mitochondrial morphological characteristics are largely similar between type I and II fibres (Figure 4), which would suggest that the number of mitochondrial profiles, rather than morphological differences, may be the primary factor that distinguishes skeletal muscle fibre types in humans. However, as we employed a two-dimensional TEM approach, we are unable to fully elucidate whether the increased number of mitochondrial profiles reflects an increased number of mitochondria or a more extensive and interconnected mitochondrial network (with increased three-dimensional foldings) - missed by the current approach. However, given that the mitochondrial network of mouse oxidative fibres is oriented in the parallel and perpendicular axis to the muscle contraction (Bleck *et al*., 2018) (while primarily perpendicular in glycolytic fibres), our results should have been capable of detecting a between-fibre morphological difference. From a mechanistic perspective, it is tempting to suggest that an optimal mitochondrial morphological range exists. By ensuring that mitochondrial size and complexity are maintained within a tight range may ensure a rapid isolation and degradation of a mitochondrion in the event of damage (Gomes *et al*., 2011; Glancy *et al*., 2017; Bleck *et al*., 2018). Furthermore, a network of individual mitochondria adjacent to each other may offer an advantage, when compared to a reticulum, to dynamically adapt to drastic changes in energy or cellular stress (i.e., exercise) (Bleck *et al*., 2018).

Mitochondrial adaptations to endurance training (Hoppeler *et al*., 1985) and inactivity (Nielsen *et al*., 2010) have been shown to occur divergently across the myofiber compartments. In the present study, we characterised mitochondrial morphological characteristics across compartments and observed an increased mitochondrial size in the SS that decreases with the intermyofibrillar depth (SS > SMF > CMF; Figure 5) in agreement with previous reports (Huertas *et al*., 2019; Castro-Sepulveda *et al*., 2020). Similarly, markers associated with mitochondrial morphological stress, such as increased circularity and roundness were elevated in the deepest CMF compartment while lowest in SS. We acknowledge that while these mitochondrial morphological differences may hold true for the studied human populations with a 2D approach, future research should employ a more comprehensive 3D approach and explore other populations such as endurance-trained or aged individuals.

### Mitochondrial remodelling with acute resistance exercise in strength athletes

Muscle adaptations are thought to stem from the molecular response to each exercise session (Perry *et al*., 2010; Viggars *et al*., 2023), which when repeated over time leads to phenotypical differences (Chapman *et al*., 2020). Our results show that a single extensive resistance exercise session in strength athletes, leads to mild changes in mitochondrial morphological variables, in the SS and to a lower extent in the SMF, associated with mitochondrial stress (Figure 6). This may indicate that resistance exercise *per se* can lead to mild degrees of mitochondrial morphological stress. Previous studies exploring mitochondrial morphological changes following different endurance exercise prescriptions (Huertas *et al*., 2019) has shown that high-intensity exercise leads to increased mitochondrial area across subsarcolemmal and intermyofibrillar mitochondria. Combined with increased complexity morphological markers, like form factor and aspect ratio, the authors suggested an exercise-induced increase in mitochondrial fusion. Conversely, we suggest that a single session of resistance exercise may induce mild mitochondrial stress (i.e., swollen mitochondria) and fragmentation (i.e., increased roundness), as we observed decreased complexity following resistance exercise - suggestive of fission. The difference between our study and that of Huertas *et al*. (2019) may be the result of a divergent mitochondrial dynamics activation. While their work suggests that endurance exercise leads to a ‘pro-fusion’ state (Huertas *et al*., 2019), our results suggest that resistance exercise may lead to a ‘pro-fission’ state. A potential explanation would have been increased mitochondrial damage, however, there were no significant changes in damaged mitochondria following resistance exercise. Future research should explore whether the mitochondrial morphological response to resistance exercise is similar across untrained and trained individuals and to which extent it is affected by other modes of exercise.

### Mitochondrial-associated transcriptional signalling following acute resistance exercise

Mitochondria adapts to changing cellular cues with the regulation of mitochondrial biogenesis and mitochondrial dynamics, which is composed of mitochondrial fusion, fission and mitophagy (Trewin *et al*., 2018). As we observed a mild mitochondrial morphological stress following resistance exercise, we interrogated the gene expression regulation of distinct mitochondrial-related pathways following resistance exercise. Using the MetaMEx database (Pillon *et al*., 2020), we did not observe any coordinated transcriptional activation of mitochondrial dynamics components, except for *MIEF2* (a fission receptor), which may suggest an increased need for mitochondrial fission. Conversely, activation of genes involved in mitochondrial biogenesis and the mitochondrial unfolded protein response (UPR^mt^) were noted. Mitochondrial biogenesis is well known to be a key pathway in exercise-induced mitochondrial remodelling (Hood *et al*., 2011), especially in response to endurance training (Granata *et al*., 2021). On the contrary, little is known regarding the role of UPR^mt^ in the exercise-induced mitochondrial remodelling (Merry & Ristow, 2016). A recent study has shown that the UPR^mt^ is upregulated in highly trained endurance athletes and to be positively associated with metabolic health (Houzelle *et al*., 2021). Therefore, we utilised a previously published RNA-seq (Chapman *et al*., 2020) of strength-trained individuals to evaluate the presence of a UPR^mt^ signature, observing an enrichment of UPR^mt^ in the resting transcriptome of strength-trained individuals (Figure 6.B), without any concurrent enrichment of mitophagy or mitochondrial dynamics genesets. Collectively, our gene expression analyses suggest that UPR^mt^ enrichment is a key signature of strength-trained individuals and that it may influence the mitochondrial phenotype associated with strength athletes. Future studies should explore these pathways at the protein level to confirm their role in the skeletal muscle phenotype of strength athletes.

## Conclusion

In summary, the present results demonstrate that strength athletes have distinct mitochondrial characteristics with decreased mitochondrial profile size, increased numeric density, increased cristae density, and increased surface-to-volume ratio compared to untrained age-matched controls. These alterations may provide strength athletes with an optimised mitochondrial network with enhanced capacity for high and rapid energy production and metabolite turnover. The present analysis also revealed that type I fibres have a higher numerical density of mitochondria than type II fibres, but with no morphological differences between the two fibre types. At the subcellular level, the SS mitochondria are bigger, and more complex across groups − with these mitochondrial characteristics decreasing with the myofiber depth. Finally, our results indicate that resistance exercise leads to signs of mild mitochondrial morphological stress, which is associated with increased gene expression of markers of mitochondrial biogenesis, fission, and UPR^mt^ − the latter representing a gene signature of strength-trained individuals.

